# The free fatty acid 2 receptor regulates the activation potency of C5a and the sensitivity to the antagonist avacopan

**DOI:** 10.1101/2025.10.28.685092

**Authors:** Zahra Khan, Neele K. Levin, Claes Dahlgren, Martina Sundqvist, Fariha Khan, Huamei Forsman, Lena Björkman

## Abstract

The complement derived neutrophil chemoattractant C5a, is a potent activator of the neutrophil superoxide anion generating NADPH oxidase. An allosteric modulator specific for the free fatty acid 2 receptor increases the activating potency but not the efficacy of C5a. The allosteric modulator also decreases the inhibitory effect of the C5a receptor antagonist avacopan, suggesting that the NADPH oxidase is activated by two different signaling pathways downstream of the receptor for C5a. While the allosteric modulator affected the C5a-mediated activation of the NADPH oxidase, the C5a-induced rise in the intracellular concentration of free calcium ions was unaffected. The C5a receptor and the free fatty acid receptor belong to the family of G protein-coupled receptors family. Our results show that the activated C5a receptors generate signals that directly activate the NADPH oxidase and allosterically modulated free fatty acid receptors which secondarily generate signals that elicit NADPH oxidase activity. This is in line with an earlier described receptor transactivation model, by which the fatty acid receptor is activated by receptor downstream signals generated by several different neutrophil receptors to which we now add the receptor for C5a. In addition, the fatty acid receptor was higher ranked than the receptor for C5a, in the neutrophil receptor hierarchy. The dual receptor trans-regulatory effects, by which the receptor for C5a activates the fatty acid receptor and by which this receptor reduces the C5a response, represent new regulatory mechanisms of importance for the NADPH oxidase activity in neutrophils.

## Introduction

The neutrophil granulocyte, the most abundant white blood cell in peripheral human blood, is of main importance for our innate immune defense system and for proper regulation of inflammatory reactions (1, 2). These cells express several cell surface receptors that regulate different cell functions such as phagocytosis of invading microbes and damaged tissue components, chemotactic recruitment of the neutrophils to infected/inflamed tissues, secretion of inflammatory mediators/proteolytic enzymes stored in the neutrophil granules and, production of reactive oxygen species (ROS) generated by a specialized electron transporting NADPH oxidase (3). Several of the neutrophil cell surface receptors belong to the family of seven transmembrane receptors also known as G protein-coupled receptors (GPCRs). Members of this receptor family are of importance for the regulation of a plethora of essential physiological processes including the basic function of inflammatory cells (4, 5). In accordance with the importance of the neutrophil GPCRs, a new receptor antagonist, avacopan, which specifically targets the C5a receptor1 (C5aR1, CD88; (6, 7)), was recently added to the growing number of available drugs directed against members of this receptor family (8–10). Two cognate C5aRs (C5aR1 and C5aR2) are expressed in human neutrophils, receptors that differ in their functional profiles; whereas C5aR1 is the primary receptor that mediates the responses induced by the agonist, C5aR2 is an atypical receptor that lacks G protein coupling, possibly with a context-dependent anti-inflammatory or immunoregulatory role (11–13). C5a is a cleavage product of complement factor C5 that is generated during complement activation (12, 14), and a well-characterized chemotactic factor mediating a balanced functional response in neutrophils upon binding to C5aR1 (15–17). The signal transduction cascade downstream of the activated C5aR1, relies on coupling to a Gα_i-_containing heterotrimeric G protein localized on the cytosolic side of the plasma membrane (18). Many neutrophil chemoattractants (but not all) that induce an increase in the intracellular concentration of free calcium ions ([Ca^2+^]_i)_, also trigger production of NADPH oxidase derived ROS upon binding to their cognate GPCR (19). This applies also to the C5a/C5aR1 ligand/receptor pair which has been described to potently activate the NADPH oxidase (20). The ROS produced by the superoxide anion (O_2-_; the precursor of all ROS) generating NADPH oxidase system are crucial for our host defense due to their capacity to kill invading pathogens (21). However, since the ROS in addition to being destructive to microbes also are toxic to host cells and tissues, activation as well as termination of the oxidase activity must be strictly regulated (19). Accordingly, blood neutrophils are in a resting (naïve) state that produce low (or no) levels of NADPH oxidase-derived ROS. The background to this is, in addition to a lack of activating agonists, that the membrane component of the NADPH oxidase (the b cytochrome) as well as several GPCRs are not present in the plasma membrane but localized in mobilizable intracellular granules/vesicles (22). However, in response to bacterial components such as lipopolysaccharides (LPS) from Gram negative bacteria (23) and inflammatory mediators such as tumor necrosis factor (TNF; (24)), neutrophils mobilize granule-stored membrane components. The increase in the plasma membrane of new GPCRs and cytochrome b, transfers neutrophils to a pre-activated/primed state characterized by an enhanced NADPH oxidase activity when triggered with certain GPCR agonists (19, 25, 26). A new priming mechanism was recently described when positive allosteric modulators (PAMs), specific for free fatty acid 2 receptor (FFA2R), were shown to play a key role for an augmentation of NADPH oxidase activity (27, 28). Accordingly, the FFA2R specific allosteric modulator Cmp58 turns not only orthosteric agonists specific for FFA2R to more potent activating ligands, but also turns several other GPCR agonists (unrelated to FFA2R) such as formyl peptides (recognized by FPRs), the β-oxidation intermediate 3-hydroxy-octanoic acid (recognized by hydroxycarboxylic acid 3 receptor; HCA_3R_) and the lipid chemoattractant LTB_4 (_recognized by BLT_1R_), into potent neutrophil activators (27, 29–31). These results violate the commonly accepted restriction that defines receptor selectivity of allosteric GPCR modulators.

In this study we have investigated C5a-induced NADPH oxidase activity in neutrophils by determining the effects of the positive allosteric modulator (PAM) Cmp58, specific for FFA2R, on the C5a-induced response. Our data confirm previous results showing that C5a alone is a strong NADPH oxidase activator (32), and this response was fully inhibited by the specific C5aR1 antagonist avacopan. The presence of the allosteric FFA2R modulator Cmp58 increased the potency of C5a but did not cause any effect on the efficacy. Moreover, Cmp58 attenuated the inhibitory effect of avacopan on the C5a-mediated response. In addition, our data show that an activation of FFA2R was accompanied by a reduced C5a response, suggesting that FFA2R is higher ranked than C5aR1 in the neutrophil receptor hierarchy.

## Material and Methods

### Chemicals

Dextran T500 and Ficoll-Paque Plus Medium used for the isolation of neutrophils were obtained from Pharmacosmos (Holbaek, Denmark) and Fischer Scientific (Gothenburg, Sweden), respectively. Bovine serum albumin (BSA), isoluminol, TNF, horseradish peroxidase (HRP), propionic acid, compound 58 (Cmp58; (S)-2-(4-chlorophenyl)-3,3-dimethyl-N-(5-phenylthiazol-2-yl)butanamide), and ATP were purchased from Sigma-Aldrich (Merck, Burlington, MA, USA). Fura-2-acetoxymethyl ester (AM) was obtained from Invitrogen by Thermo Fisher Scientific (Gothenburg, Sweden) and CATPB ((S)-3-(2-(3-chlorophenyl)acetamido)-4-(4-(trifluoromethyl)phenyl) butanoic acid) was from Tocris (Bristol, UK). Recombinant human C5a (rhC5a) and AZ1729 was purchased from R&D Systems (Minneapolis, MN, USA) and avacopan was purchased from MedChemExpress (Princeton, NJ, USA). Stock solutions were prepared in DMSO, and working solutions were prepared in Krebs-Ringer glucose phosphate buffer (KRG, 120 mM NaCl, 4.9 mM KCl, 1.7 mM KH_2P_O_4,_ 8.3 mM Na_2H_PO_4,_ 1.5 mM MgSO_4,_ 10 mM glucose, and 1 mM CaCl_2 i_n dH_2O_, pH 7.3).

### Ethical statement

The present study comprises buffy coats, which were obtained from healthy human blood donors at the blood bank of Sahlgrenska University Hospital in Gothenburg, Sweden. All the buffy coats were obtained anonymously and, therefore, no ethical approval was required according to the Swedish legislation section code 4§ 3p SFS 2003:460 (Law on Ethical Testing of Research Relating to People).

### Isolation of human neutrophils

Human neutrophils were isolated from buffy coats by dextran sedimentation and Ficoll-Paque gradient centrifugation as described earlier (33). After separation, contaminating red blood cells were lysed by hypotonic lysis and the remaining cells were washed and resuspended (1 x 10^7^ cells/mL) in KRG and kept on ice until used in further assays on the same day as the isolation. The purity of neutrophils was determined in an automatic analyzer (Sysmex KX-21 N Hematology Analyzer, Sysmex Corporation) and routinely contained ≥ 90% neutrophils. To amplify the activation signals of the neutrophil NADPH oxidase, freshly isolated neutrophils were primed with TNF (10 ng/mL; 1 x 10^6^ cells/mL) at 37°C for 20 minutes in a water bath and then stored on ice until use.

### Measuring NADPH oxidase activity in neutrophils

An isoluminol-enhanced chemiluminescence (CL) system was used to measure the production of superoxide anions (O_2-_) generated by the neutrophil electron transporting NADPH oxidase as described earlier (34, 35). Briefly, all the measurements were performed using a six-channel Biolumat LB 9505 (Berthold Co., Wildbad, Germany). Disposable polypropylene tubes (4 mL) were used, containing a 900 µL reaction mixture comprising 1 x 10^5^ neutrophils, isoluminol (2 x 10^-5^ M) and HRP (4 units/mL). The tubes were incubated for 5 min at 37°C before adding the activating ligand (100 µL), and the light emission was recorded continuously. To determine the effects of receptor-specific antagonists and allosteric modulators, these ligands were added to the reaction mixture 1-5 min before agonist stimulation. Controls were run in parallel for comparison. The O_2-_ production, recorded continuously over time, was expressed in Mega counts per minute (Mcpm) and peak activities were used for comparisons of the NADPH oxidase activities.

### Measurement of the cytosolic concentration of free calcium ions

An increase in the intracellular concentration of free calcium ions ([Ca^2+^]_i)_ in neutrophils was measured by utilizing the Ca^2+^ sensitive dye Fura 2-AM. Neutrophils were Fura 2-AM loaded and analyzed for an agonist-mediated rise in [Ca^2+^]_i b_y fluorescence spectrophotometry as described previously (36). In brief, isolated neutrophils (2 × 10^7^ cells/mL in KRG containing 0.1% BSA) were loaded with Fura 2-AM (2 µg/mL, 30 min at room temperature in darkness) and thereafter washed twice before resuspended to a concentration of 2 × 10^7^ neutrophils/mL in ice cold KRG. The Fura 2-AM stained neutrophils, were kept on ice in darkness until measurements of [Ca^2+^]_i o_n a Perkin Elmer fluorescence spectrometer (LS50B), with excitation wavelengths of 340 nm and 380 nm, an emission wavelength of 509 nm, and slit widths of 5 nm and 10 nm, respectively. For the measurements, disposable polystyrene cuvettes (4.2 mL) containing 2.475 mL reaction mixture (2 × 10^6^ Fura 2-AM stained neutrophils/ml in KRG) were first equilibrated at 37°C for ten minutes, prior addition of an agonist (25 µL). The transient increase in [Ca^2+^]_i o_ver time is shown as the ratio of the fluorescence intensities (340 nm: 380 nm) determined.

### Statistical data analysis

The raw data obtained were analyzed for statistically significant differences either by repeated measures one-way analysis of variance (ANOVA) or mixed-effects analysis followed by Šίdák’s multiple comparison test, or unpaired or paired Student’s t-test, using GraphPad Prism 8.4.2 (GraphPad Software, San Diego, CA, USA). The results obtained from the data are presented as a mean with a standard error of the mean (SEM). A *p*-value ≤ 0.05 was considered a statistically significant difference (denoted as * *p* ≤ 0.05, ** *p* ≤ 0.01, *** *p* ≤ 0.001. The number of repeats performed independently on neutrophils isolated from different individuals (n) and the specific statistical test used for each figure are described in the figure legends.

## Results

### The complement component C5a is a potent NADPH oxidase activating agonist and the response is further increased by the priming cytokine TNF

Neutrophils express an electron-transporting enzyme system (the NADPH oxidase; (21)) that when activated, transfers electrons over the plasma membrane from NADPH in the neutrophil cytosol to extracellular localized molecular oxygen that becomes reduced to superoxide anions (O_2-_). The chemoattractant C5a, a peptide fragment generated from the complement component C5 during complement activation (14), is a potent NADPH oxidase activating agonist in neutrophils (Fig 1A).

**Figure 1.**
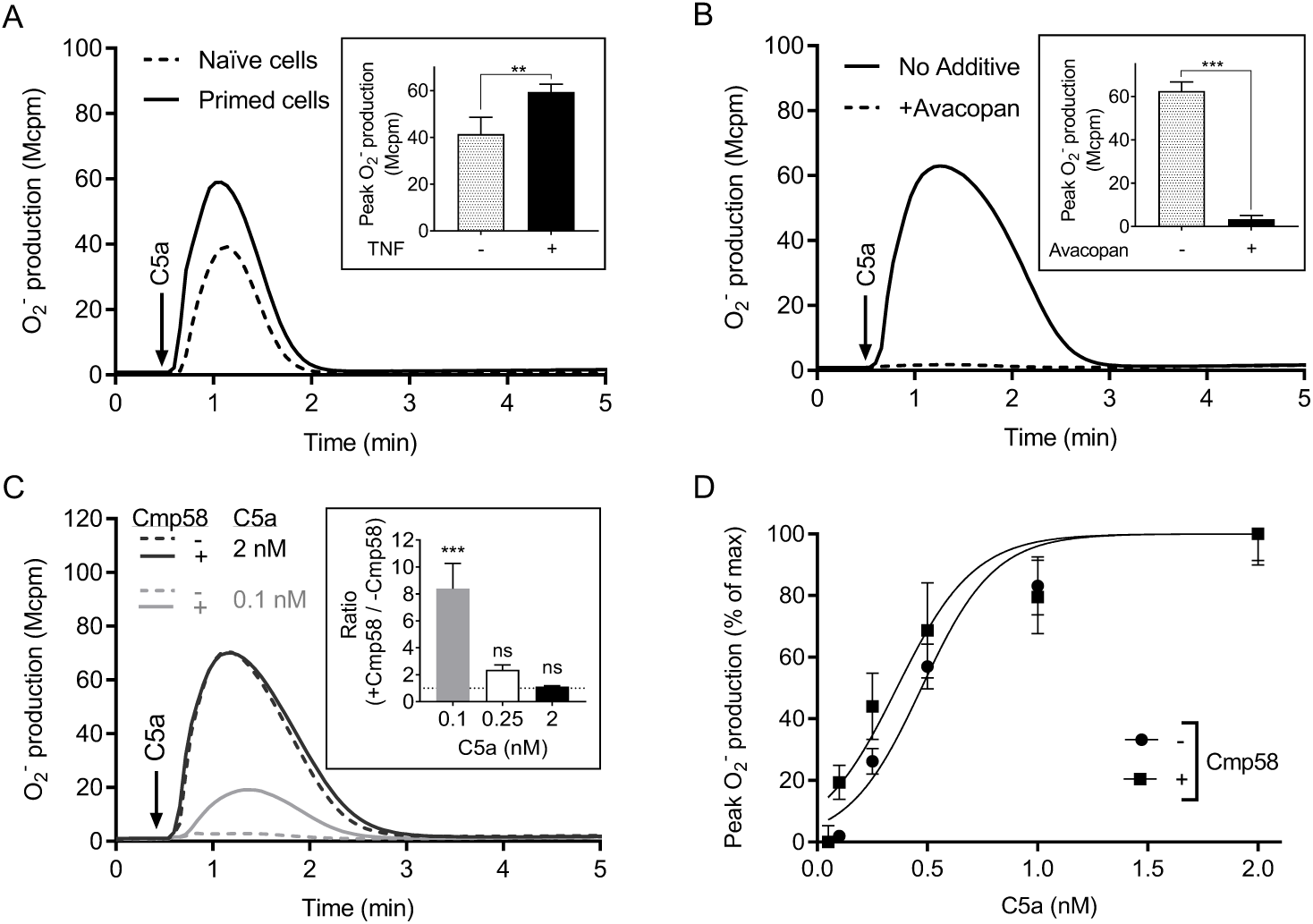
TNF primes neutrophils and an allosteric FFA2R modulator turns C5a into a more potent activator of the NADPH oxidase. For priming, neutrophils were incubated with TNF (10 ng/mL) at 37°C for 20 min and then stored on ice until used. The NADPH oxidase activity (O_2-_ production) induced by C5a was measured continuously. **(A)** The response induced in naïve (non-primed; dashed line) and TNF primed neutrophils (solid line) by C5a (2 nM, added at the time point marked with an arrow) was measured. **Inset:** The priming effect of TNF on the NADPH oxidase activity expressed as peak values (Mcpm; mean ± SEM) of O_2-_ production induced by C5a in naïve (n = 5) and TNF-primed neutrophils (n = 14). **(B)** Inhibition of the C5a-induced response by the specific C5aR1 antagonist avacopan. TNF-primed neutrophils were incubated without (solid line) or with avacopan (50 nM, dashed line) for 5 min before addition of C5a (2 nM; added at the time point marked with an arrow) and determination of the NADPH oxidase activity. **Inset:** The inhibitory effect of avacopan on the neutrophil NADPH oxidase activity expressed as peak values (Mcpm) of O_2-_ production induced by C5a in absence and presence of the antagonist, respectively (mean ± SEM, n = 6). **(C)** Effects of the allosteric FFA2R modulator Cmp58 on the response induced by different concentrations of C5a in TNF-primed neutrophils. Neutrophils were incubated without (dashed lines) or with Cmp58 (1 µM, solid lines) for 5 min before addition of C5a (0.1 nM, grey lines, or 2 nM, black lines; added at the time point marked with an arrow) and determination of the NADPH oxidase activity. **Inset:** The effect of Cmp58 on the NADPH oxidase activity expressed as fold increase of O_2-_ production induced by different C5a concentrations (0.1, 0.25 or 2 nM) in the absence and presence of Cmp58, respectively (mean ± SEM, n = 6). The horizontal dotted line in the bar graph represents a ratio of 1. **(D)** TNF-primed neutrophils were pre-incubated with and without Cmp58 (1 µM) for 5 min and activated with different concentrations of C5a as indicated. Superoxide production was recorded continuously and expressed as the peak value obtained in terms of percent of the activity induced by C5a alone (2 nM, mean ± SEM, n = 3). One representative experiment is shown in each subset (**A, B,** and **C**). Statistically significant differences in the insets were evaluated by an unpaired Student’s *t*-test (**A**), a paired Student’s *t*-test (**B**), or a repeated measures one-way ANOVA followed by Šίdák’s multiple comparison (**C**) and are denoted as ** (*p* ≤ 0.01), *** (*p* ≤ 0.001), and ns = not significant.

TNF is a potent and commonly used neutrophil priming agent (37, 38) that lacks direct effect on the NADPH oxidase but amplifies the neutrophil response induced by several different G protein-coupled receptors (19). In accordance with this, also the C5a-induced response was primed, meaning that the response was increased in the neutrophils pre-treated with TNF, as compared to the response induced in naïve cells (Fig 1A). In accordance with the involvement of the neutrophil C5aR1 in the C5a-induced response, the presence of the receptor-selective antagonist avacopan (6) fully inhibited the response induced by C5a (Fig 1B).

The response induced by C5a was of the same magnitude as that induced by the prototypic neutrophil activating peptide fMLF, but whereas the EC_50-_value of fMLF is ≈ 30nM (39, 40) the value of C5a is ≈ 0.5 nM (Fig 1D).

Based on previous published data (27, 30, 31), showing that TNF priming is of importance in a novel receptor transactivation mechanism involving the free fatty acid 2 receptor (FFA2R), TNF-primed neutrophils were used in the experiments described below, designed to determine FFA2R dependent trans-regulating mechanisms in neutrophils activated by C5a.

### The allosteric FFA2R modulator Cmp58 increases the potency but not the efficacy of C5a

Cmp58, an earlier described positive allosteric FFA2R modulator, turns not only non-activating orthosteric FFA2R agonists such as the short chain free fatty acid propionate into NADPH oxidase activating ligands, but also other non- or low-activating agonists recognized by other GPCRs expressed in neutrophils, are turned into potent neutrophil activating ligands (27). The precise mechanism for this activation has not yet been disclosed but an attractive model for how signals generated by one neutrophil GPCR transactivate the allosterically modulated FFA2R from the cytoplasmic side of the plasma membrane has been presented (19). To determine if Cmp58 also potentiates the response induced in neutrophils by low- or non-activating concentration of C5a, the allosteric FFA2R modulator was included in the system designed to measure NADPH oxidase activity. The results obtained using three different C5a concentrations to activate the neutrophils illustrates the effect of Cmp58; with a low (0.25 nM) or a very low (0.1 nM) concentration of C5a, the response was substantially increased in the presence of Cmp58 (Fig 1C and D). Cmp58 did, however, not affect the magnitude of the response induced by a high concentration of C5a (2 nM) that, on its own, fully activates the NADPH oxidase (Fig 1C and D).

### Cmp58 changes the inhibitory effect of the C5aR1 specific antagonist avacopan

When applying the suggested two-receptor transactivation model (19), on the Cmp58-primed C5a response, a C5aR1 specific antagonist alone, but also an FFA2R specific antagonist alone, should inhibit the response. This inhibition pattern agrees with the results obtained when neutrophils were activated by a very low concentration (0.1 nM) of C5a, as this response was separately inhibited both by the C5aR1 antagonist avacopan and the FFA2R antagonist CATPB (Fig 2A). The fact that both C5aR1 and FFA2R participate in the C5a-induced activation of the neutrophil NADPH oxidase fully support a receptor transactivation mechanism by which the signals generated by the agonist occupied C5aR1 activate the allosterically modulated FFA2R to activate the NADPH oxidase. This is also supported by the partial inhibition of the Cmp58 amplified response induced by 0.25 nM C5a (Fig 2B).

**Figure 2.**
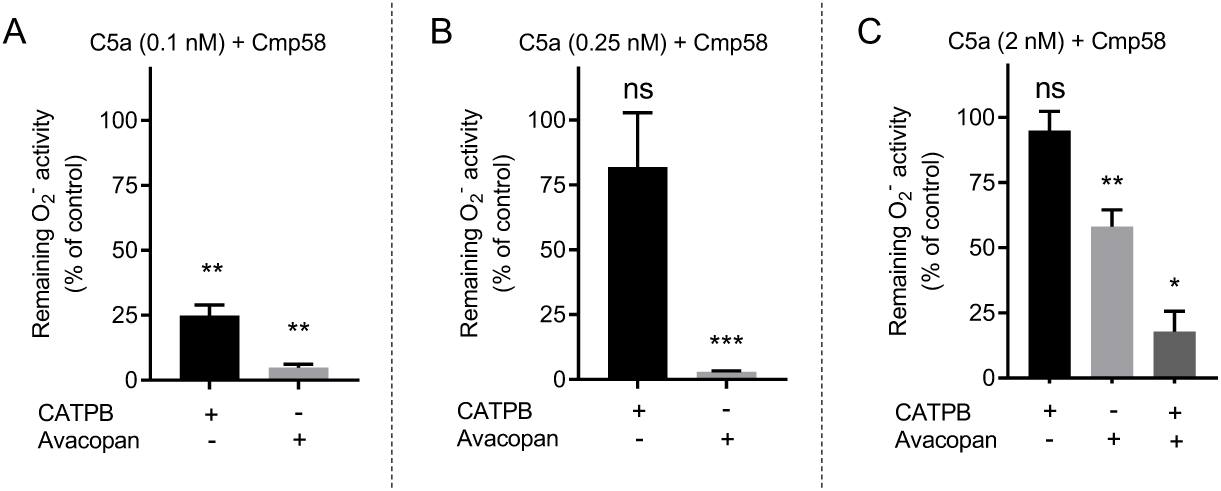
Antagonists selective for FFA2R and C5aR1, respectively, have inhibiting profiles beyond the expected receptor specificity. TNF-primed neutrophils were incubated with Cmp58 (1 µM; 5 min at 37°C) and to determine the inhibitory effects of receptor specific antagonist, these cells were incubated without or with an antagonist (CATPB, 100 nM; avacopan, 50 nM) and the O_2-_ production was measured continuously following an activation by different concentrations of C5a. **(A, B, C)** Inhibition by the antagonists (CATPB and avacopan) of the response in neutrophils incubated with Cmp58 and induced by C5a, at different concentrations, i.e., 0.1 nM (**A**; n = 6), 0.25 nM (**B**; n=7), and 2 nM (**C**; n=3-8), respectively. Inhibition is expressed as the remaining activity (peak value in percent) of neutrophils after activation with C5a in the presence of the respective antagonist. Bar graphs are presented as mean ± SEM. The statistical determinations are based on the difference between the C5a response without and with the antagonists. Statistically significant differences were evaluated by a repeated measures one-way ANOVA followed by Šίdák’s multiple comparison (**A-B**) or by a mixed effects analysis followed by Šίdák’s multiple comparison (**C**) and are denoted as * (*p* ≤ 0.05), ** (*p* ≤0.01), *** (*p* ≤ 0.001), and ns = not significant.

The response induced by a high concentration of C5a (2 nM) is fully inhibited by avacopan (see Fig 1B), and this response is not increased by Cmp58 (Fig 1C and D). However, in contrast to the potent inhibitory effect by avacopan on the response induced by C5a alone (Fig 1B), the presence of Cmp58 clearly reduced the inhibitory effect of avacopan (Fig 2C). Even though Cmp58 alone lacked effect on the 2 nM C5a-induced response that also was insensitive to the FFA2R antagonist CATPB, the inhibitory profile of avacopan was restored when CATPB was added to the measuring system (Fig 2C). These results suggest that two different signaling pathways are activated by the agonist occupied C5aR1 – one pathway requires that many receptors are occupied and this signaling pathway directly activates the neutrophil NADPH oxidase, and another signaling pathway is initiated also when very few receptors are occupied, and this pathway activates the allosterically modulated FFA2R.

### Neutrophil NADPH oxidase activity induced when the order was reversed, by which the allosteric modulator and the activating agonist were added to the neutrophils

One of the basic characteristics of the relation between a receptor specific PAM and an activating agonist recognized by the same receptor, is that the order by which the two ligands are added to the receptor-expressing cell can be reversed. In contrast, we have in several studies shown that this type of reciprocity is not obtained when FFA2R is transactivated by signals generated by another GPCR (19, 27, 30, 31, 41). The characteristic pattern of ligand reversibility is illustrated by the relation between Cmp58 and propionate. That is, the NADPH oxidase is activated in neutrophils stimulated with propionate in the presence of the allosteric FFA2R modulator Cmp58 (Fig 3A) and the NADPH oxidase is activated also when the order by which the two FFA2R ligands are added to the cells are reversed (Fig 3B). No reversibility was, however, seen when C5a replaced propionate; although a similar NADPH oxidase activity as that with propionate was obtained in neutrophils activated by C5a (Fig 3A), no NADPH oxidase activity was obtained when the order by which Cmp58 and C5a was added to the neutrophils was reversed (Fig 3B).

**Figure 3.**
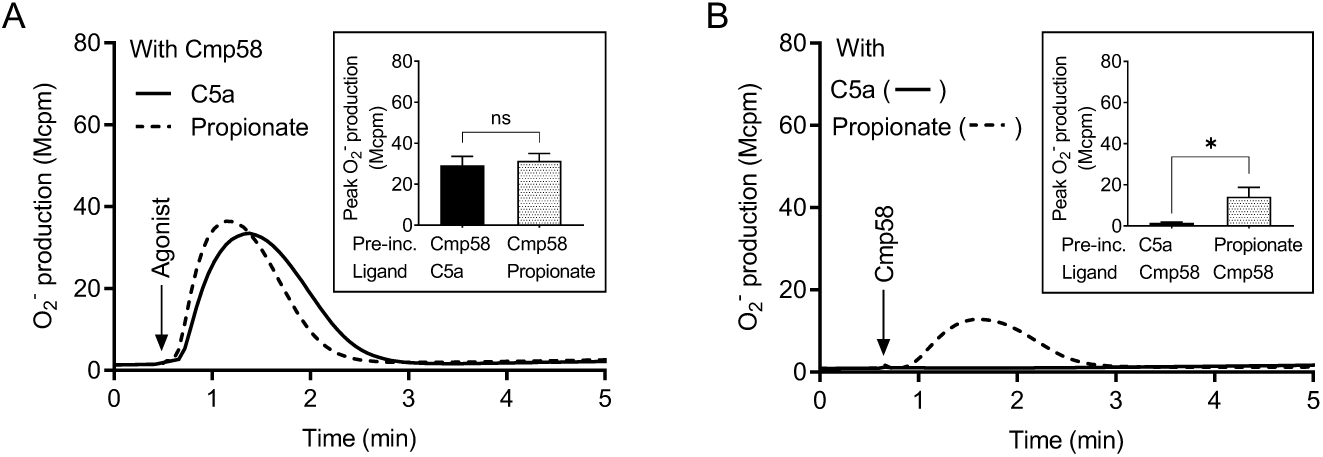
Differential NADPH oxidase activity triggered by Cmp58 in neutrophils pretreated with C5a or propionate. **(A)** TNF-primed neutrophils pre-incubated with Cmp58 (1 µM, 5 min at 37°C) were activated by C5a (0.1 nM, solid line) or propionate (25 µM, dashed line). The result obtained in one representative experiment is shown. **Inset:** The NADPH oxidase induced by the ligands C5a and propionate, in neutrophils pre-incubated (abbreviation Pre-inc.) with Cmp58, expressed as the peak values of O_2-_ production (mean ± SEM, n = 5). **(B)** TNF-primed neutrophils incubated for 5 min at 37°C with non-activating concentrations of C5a (solid line; 0.1 nM) or propionate (dashed line; 25 µM) were activated by Cmp58 (1 µM). The result obtained in one representative experiment is shown. **Inset:** The NADPH oxidase induced by Cmp58, in neutrophils preincubated (abbreviation Pre-inc.) with C5a and propionate, respectively, expressed as the peak values of O_2-_ production (mean ± SEM, n = 5). Statistically significant differences in the insets were evaluated by a paired Student’s *t*-test and are denoted as * (*p* ≤ 0.05) and ns = not significant.

### Cmp58 does not affect the rise in the cytosolic concentration of free calcium ions ([Ca^2+^]_i)_ in C5a activated neutrophils

A model described for receptor crosstalk synergy between the C5aR1 and the purinergic receptor that recognize UDP has been suggested to involve the G protein mediated rise in the cytosolic concentration of free calcium ions ([Ca^2+^]_i)_ (42). To determine a potential involvement of [Ca^2+^]_i i_n the crosstalk between C5aR1 and FFA2R, both being Gα_i-_coupled GPCRs, we determined the effects of the allosteric modulator Cmp58 on the transient rise in [Ca^2+^]_i m_ediated by different concentrations of C5a. In agreement with earlier findings, low non-activating concentrations of propionate (25 µM) induced a rise in [Ca^2+^]_i w_hen FFA2R was allosterically modulated by Cmp58, an experiment serving as a positive control (Fig 4). C5a triggered an agonist concentration-dependent rise in [Ca^2+^]_i i_n neutrophils, but this response was not affected by Cmp58 irrespective of the C5a concentration (Fig 4). This shows that signaling in the presence of Cmp58 is biased in a way that it remains inert in the C5aR1-mediated rise in [Ca^2+^]_i,_ but at the same time increases the production of superoxide anions in neutrophils activated by low concentrations of C5a (Fig1 C-D, 2, and 4).

**Figure 4.**
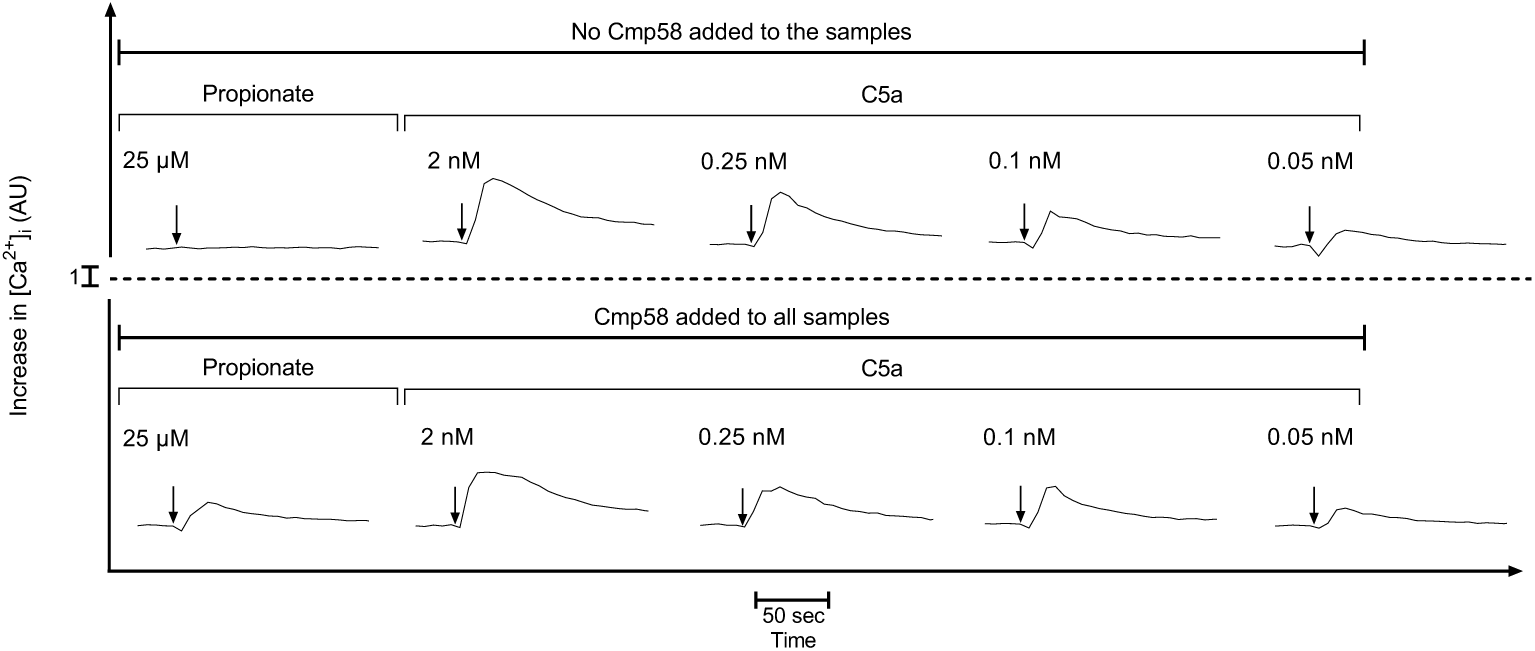
C5a triggers a transient increase in the cytosolic concentration of free calcium ions ([Ca^2+^]_i)_ in neutrophils, which is not affected by the allosteric FFA2R modulator Cmp58. Fura-2 loaded neutrophils were activated by propionate (25 µM) or different concentrations of C5a (2, 0.25, 0.1, and 0.05 nM) in the absence (upper panel) or presence (lower panel) of the allosteric FFA2R modulator Cmp58 (1 µM, pre-incubated with the cells for 10 min at 37°C prior addition of the agonist). Results obtained are shown as one representative experiment out of 3. The arrows show the time point for the addition of the agonists to the Fura-2 labelled neutrophils.

### Inhibition of the C5a-induced NADPH oxidase activity in neutrophils first activated by an FFA2R activating ligand

The three ligands (propionate, ATP and AZ1729) used to activate FFA2R have alone no NADPH oxidase activating capacity. The allosteric FFA2R modulator Cmp58 turns, however, these ligands into potent NADPH oxidase activating ligands that initiate this FFA2R dependent signaling by different mechanisms; i) the orthosteric agonist propionate activates the allosterically modulated FFA2R to produce O_2-_ (Fig 5A) a response coupled to a rise in [Ca^2+^]_i,_ ii) ATP, an agonist recognized by the Gα_q-_coupled purinergic receptor P2Y_2R_, transactivates the allosterically modulated FFA2R to produce O_2-_, (Fig 5B), and iii) also AZ1729, a positive allosteric FFA2R modulator that binds to a unique allosteric receptor site activates the Cmp58 modulated FFA2R to produce O_2-_ (Fig 5C) but with a signaling profile that differs from that used by propionate (43). The response induced by C5a, once the FFA2R/Cmp58 dependent response was terminated, was diminished (desensitized) irrespectively of which of the three ligands that was used to activate FFA2R signaling (Fig 5A-C). Notably, the inhibitory response varied depending on how FFA2R was activated, being most pronounced in neutrophils initially activated with AZ1729. Taken together, these data suggest that an activation of FFA2R partly desensitizes C5aR1.

**Figure 5.**
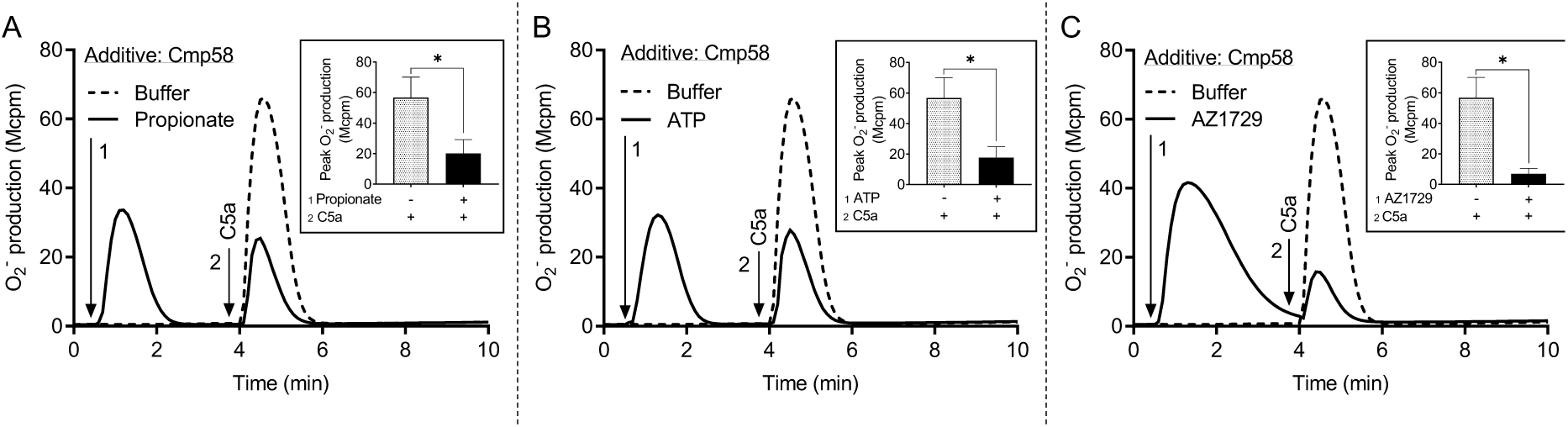
Activation of FFA2R inhibits (heterologously desensitizes) the neutrophil response induced by C5a. TNF-primed neutrophils incubated with Cmp58 (1 µM), were either left in a resting state (dashed lines) or activated by an FFA2R activating/transactivating ligand (solid lines) and the O_2-_ production was measured continuously. When the response induced by the FFA2R activating agonist was terminated, the two cell samples were activated by an addition of C5a (2 nM). The results obtained in one representative experiment is shown together with an inset showing the peak activities induced by C5a (mean ± SEM, n = 3). **(A)** Propionate (25 µM) was used as the FFA2R activating ligand (the time point for addition is marked by arrow 1). The time point for addition of C5a to the cells is marked by arrow 2. **(B)** ATP (25 µM) was used as the FFA2R activating ligand (the time point for addition is marked by arrow 1). The time point for addition of C5a to the cells is marked by arrow 2. **(C)** AZ1729 (1 µM) was used as the FFA2R activating ligand (the time point for addition is marked by arrow 1). The time point for addition of C5a to the cells is marked by arrow 2. Statistically significant differences in the insets were evaluated by a paired Student’s *t*-test and are denoted as * (*p* ≤ 0.05).

### Homologous desensitization of FFA2R mediated by an activation of the C5aR1

Sequential receptor activation to determine the hierarchical relationship between FFA2R and C5aR1, showed that the C5a response in neutrophils first stimulated with an FFA2R activating ligand, was inhibited (heterologous desensitized). This suggests that FFA2R has a higher rank than C5aR1 (Fig 5). When the receptor activation order was reversed, the response induced by an FFA2R activation ligand in neutrophils first activated by C5a, was also substantially reduced. This inhibition was obtained irrespectively if FFA2R activation was mediated by the orthosteric FFA2R agonist propionate (Fig 6A), the transactivating P2Y_2R_ agonist ATP (Fig 6B), or by the allosteric FFA2R modulator AZ1729 (Fig 6C). More important, however, is that the inhibitory effect coupled to C5a-induced activation required the presence of the allosteric FFA2R modulator Cmp58 during activation by C5a. No inhibition of the response induced by propionate (Fig 6D), ATP (Fig 6E) or AZ1729 (Fig 6F) was obtained when Cmp58 was added once the C5a-induced response had been terminated (Fig 6D-F). Taken together, these data suggest that the inhibition mediated when the C5aR1 is activated in the presence of Cmp58 is homologous rather than heterologous, and the result of a Cmp58 dependent effect of a C5aR1-mediated activation/desensitization of FFA2R.

**Figure 6.**
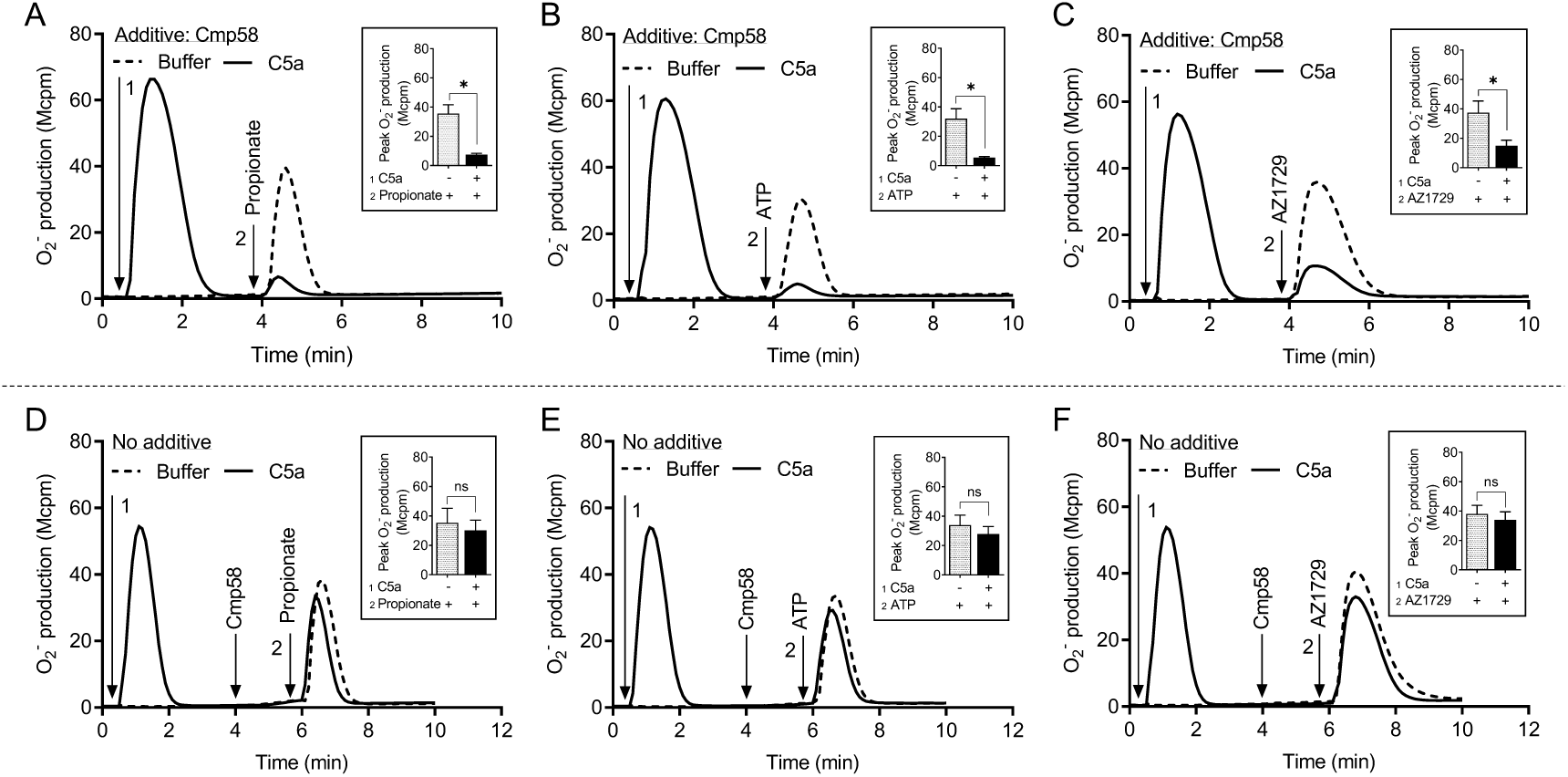
C5a-induced desensitization in Cmp58-sensitized neutrophils. **(A, B, C)** TNF-primed neutrophils incubated with Cmp58 (1 µM, 5 min at 37°C) were either left in a resting state (dashed lines) or activated by C5a (2 nM, solid line; the time point for addition is marked by arrow 1), and the O_2-_ production was measured continuously. When the response, induced by the C5aR1 activating ligand was terminated, the two cell samples were activated by the addition of **(A)** propionate (25 µM, an FFA2R activating ligand), **(B)** ATP (50 µM, a P2Y_2R_ activating ligand), or **(C)** AZ1729 (1 µM, an FFA2R activating ligand), and the time point for their addition is marked by arrow 2 **(D, E, F)**. The experimental setup differs from that described above, in that Cmp58 (1 µM), was added first when the response induced by C5a (time for addition marked by arrow 1), had settled. The neutrophils were then activated with either propionate (**D**; 25 µM), ATP (**E**; 50 µM), or AZ1729 (**F**; 1 µM). The time point for addition of the FFA2R activating/transactivating ligand is marked by arrow 2, and one representative experiment is depicted, and the results obtained are shown together with an inset showing the peak activities induced by the FFA2R activating/transactivating ligands (mean ± SEM, n = 3). Statistically significant differences in the insets were evaluated by a paired Student’s *t*-test and are denoted as * (*p* ≤ 0.05) and ns = not significant.

## Discussion

The endogenous inflammatory mediator C5a, a 74 amino acids long cleavage product derived from complement component C5 during activation of the complement system (44) and recognized by the G protein-coupled receptor (GPCR) C5aR1, is a potent neutrophil activating ligand (45). Although the exact receptor downstream signals that are triggered and regulate the different functional responses mediated upon C5a binding to C5aR1 are not known in detail, it is clear that C5a is a potent neutrophil chemoattractant, inducer of the PLC dependent rise in [Ca^2+^]_i a_nd activation of the superoxide anion (O_2-_) producing NADPH oxidase (46). In this study we show that in addition to the direct activation of the NADPH oxidase mediated by C5a, the presence of an FFA2R specific positive allosteric modulator (Cmp58) augments the response induced by low- and non-activating concentrations of C5a. Our data demonstrate that the Cmp58-amplified response to C5a was inhibited not only by the C5aR1-specific antagonist avacopan, but also by the FFA2R-specific antagonist CATPB, suggesting that this NADPH oxidase activation was achieved through a receptor transactivation mechanism, initiated by signals generated downstream of the C5a-occupied C5aR1. These results are consistent with a receptor transactivation model in which intracellular signals generated by the C5aR1 (as well as by several other GPCRs; see (27, 30, 31)) activate the allosterically modulated FFA2Rs from the cytosolic side of the plasma membrane. The signals generated by this receptor transactivation mechanism recruit the cytosolic oxidase components to the plasma membrane where the NADPH oxidase is assembled and activated. Since C5a alone has the capacity to activate the neutrophil NADPH oxidase, we suggest a two-signal model for how signals generated by the agonist occupied C5aR1 activate the NADPH oxidase (described in Fig 7). This two-signal model is supported by our data showing that the C5aR1 antagonist avacopan (50 nM) fully inhibits the response induced by 2 nM C5a alone. However, although Cmp58 had no additive effect on this C5a-induced response, this response was only partly inhibited by avacopan. Notably, the inhibitory effect was restored by the addition of the FFA2R antagonist CATPB, which blocks the Cmp58 induced sensitization to the transactivating signal (Fig 2). These findings suggest that only a small fraction of the C5aR1s need to be occupied to transactivate the allosterically modulated FFA2Rs, a suggestion supported by the data showing that Cmp58 reduced the inhibitory effect of avacopan when 2 nM C5a was used to activate the neutrophils, and that the inhibitory effect of the C5aR1 antagonist was restored when CATPB blocked the action of Cmp58. As mentioned, the receptor transactivation initiated by the C5a/C5aR1 agonist/receptor complex is in line with earlier published data showing that the NADPH oxidase response induced by several other GPCR agonists are also amplified through a transactivation of FFA2R. These agonists include ATP, PAF, LTB_4 a_nd non-activating concentrations of peptides with an fMet in the N-terminus (19). Based on these results, it is time to re-evaluate the prevailing paradigm of allosteric receptor modulators, which assumes that PAMs (positive allosteric receptor modulators), solely affect the response by orthosteric agonists acting on the receptor that binds the allosteric modulator (47–50).

**Figure 7.**
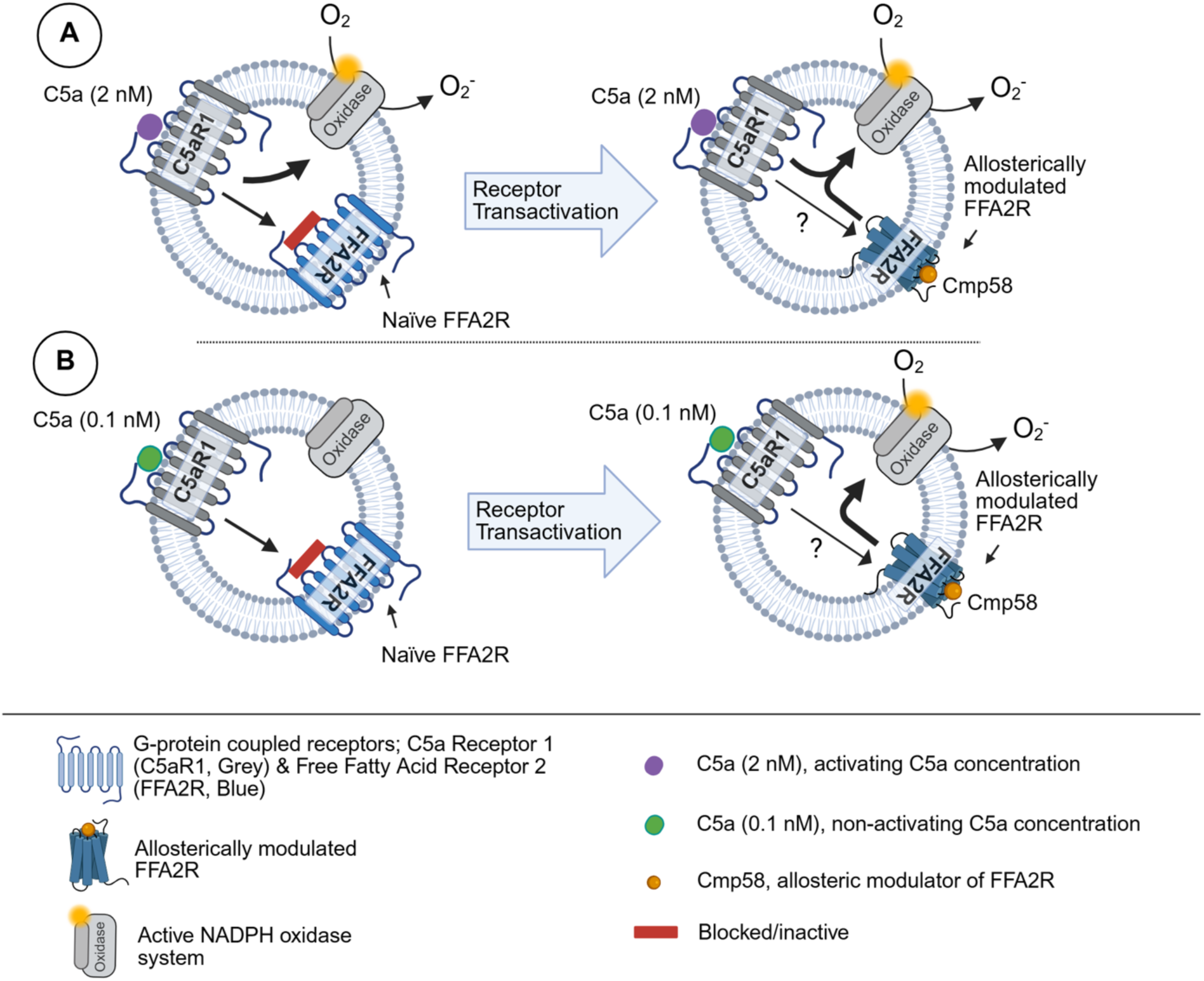
A proposed model for how the neutrophil NADPH oxidase is turned to an active state by the C5aR1 agonist C5a in the absence and presence of the allosteric FFA2R modulator Cmp58 (A) Left: The neutrophil response induced by a 2 nM concentration of C5a. In the absence of Cmp58, the signals generated by the activated C5aR1 directly activate the O_2-_-generating NADPH oxidase and have also the capacity to transactivate FFA2R, but the naïve fatty acid receptor is not receptive to these signals. **Right:** In the presence of Cmp58, the signals generated by the activated C5aR1 directly activate the O_2-_-generating oxidase and transactivate the allosterically modulated FFA2R; the transactivated FFA2R does not directly potentiate the NADPH oxidase response but reduces the inhibitory effect on this response, of the C5aR1 specific antagonist avacopan. **(B) Left:** The neutrophil response induced by a 0.1 nM concentration of C5a. In the absence of Cmp58, the signals generated by the activated C5aR1 have no NADPH oxidase activating effect. The signals have, however, the capacity to transactivate FFA2R, but the naïve fatty acid receptor is not receptive to these signals. **Right:** In the presence of Cmp58, the signals generated by the activated C5aR1 activate the O_2-_-generating NADPH oxidase, and this activation is totally dependent of the signals generated by the allosterically modulated FFA2R, a receptor made receptive to the transactivating signals generated by C5aR1.

Human neutrophils express multiple GPCRs that recognize and respond to signaling molecules belonging to different chemical classes including peptides/proteins, short chain fatty acids, lipids, nucleotides and small compounds. These GPCRs are not all equal in their functional impact but follow a receptor hierarchy, where some downstream signals dominate over others in directing neutrophil behavior (51). The proposed mechanisms behind this hierarchy include heterologous receptor desensitization by signals from a high-priority GPCR that phosphorylate a lower priority GPCR. An illustrative example of this phenomenon is the dominance of the high-ranked FPRs, which upon activation, desensitize the lower ranked IL8 receptors (CXCR1/2; (52)). C5aR1 is also regarded as a hierarchically strong receptor, sharing the top-position (or near the top position; (51)) in this hierarchy and its ability to override signals from GPCRs activated by other inflammatory mediators may serve to ensure an efficient and prioritized immune reaction. In this study, we have investigated the hierarchical relationship between the two receptors FFA2R and C5aR1 by sequentially stimulating the cells with C5a and activating ligands recognized by FFA2R. The response induced in neutrophils that were first activated with C5a was inhibited (partly desensitized) when these cells were activated by propionate, that was added once the initial C5a-induced response had subsided. This desensitization was, however, entirely dependent on the presence of the allosteric FFA2R modulator Cmp58 during the C5a-induced activation. The fact that the presence of Cmp58 is required for the C5a-induced inhibition of the FFA2R response, suggests that FFA2R is activated by signals generated by C5aR1 and that the desensitization should be classified as homologous rather than heterologous. We have previously shown that the allosterically modulated FFA2R can be activated without the involvement of any orthosteric agonist; i) FFA2R is activated by another FFA2R-specific allosteric modulator (AZ1729; (43)) that binds to a different allosteric binding site than Cmp58, and ii) FFA2R is activated by signals generated by the ATP receptor P2Y_2R_, and the signals generated by P2Y_2R_ activate FFA2R from the inside of the neutrophil plasma membrane (30). The hierarchical relationship between FFA2R and C5aR1 was maintained when the cells were sequentially stimulated with C5a, also when propionate was replaced by AZ1729 and ATP, respectively. Also, this desensitization was entirely dependent on the presence of the allosteric modulator Cmp58 when C5a was used to activate the cells. These results fully support the proposal that the desensitization is homologous and not dependent on the FFA2R agonist used to activate the desensitized neutrophils. The relationship between the two receptors was changed when the order of the activating ligands was reversed, that is, when C5a was used as the activating agonist in the second step of this sequential activation. Under these conditions, the C5a-induced NADPH oxidase activity was reduced in neutrophils that were first activated with an FFA2R-activating ligand, and the inhibitory effect was largest when AZ1729 was used as the initiating (desensitizing) ligand. The results show that C5aR1 is partially desensitized (heterologously) by FFA2R and that in neutrophils, FFA2R has a higher rank than C5aR1.

As mentioned, several different neutrophil GPCRs have the capacity to activate the NADPH oxidase through receptor transactivation, but the precise mechanism by which signals generated by these receptors (now including also C5aR1), activate FFA2Rs, has not yet been disclosed. Earlier published data suggest, however, that the transactivating signals are generated downstream of the G protein to which the activated receptor is coupled. Despite signaling differences between Gα_i (_i.e., FPRs, and BLT_1R_) and Gα_q (_i.e., PAFR and P2Y_2R_) coupled GPCRs, both activate the PLC dependent transient rise in [Ca^2+^]_i._ (19). This PLC-PIP_2-_IP_3 d_ependent signaling pathway might, thus, be part of the transactivation mechanism but such an activation mechanism requires that the allosteric FFA2R modulator transfers FFA2R to a state that is directly or indirectly activated by a rise in [Ca^2+^]_i._ This mechanistic model has gained support by data showing that such an activation of the NADPH oxidase is achieved when a rise in [Ca^2+^]_i i_s induced by the Ca^2+^-selective ionophore ionomycin or by an inhibition of the Ca^2+^-transporting ATPase present in the endoplasmic reticulum (41). Taken together, the NADPH oxidase response induced by the potent endogenous activator C5a, is increased by Cmp58, an allosteric modulator of FFA2R; the activation potency is increased without any effect on the efficacy of C5a. These results are in line with an earlier described receptor transactivation (crosstalk) model, in which FFA2R is activated by receptor downstream signals from multiple neutrophil GPCRs, now including C5aR1. Our data show that Cmp58 reduces the inhibitory effect of the C5aR1 antagonist avacopan, an effect that was reversed by the FFA2R antagonist CATPB. We conclude that the NADPH oxidase can be activated by two distinct C5aR1-dependent signaling pathways; one direct and one through transactivation of the allosterically modulated FFA2Rs that secondarily generates signals which elicit NADPH oxidase activity (Fig 7). The receptor trans-regulation, by which the C5aR1 activates FFA2R and by which the activated FFA2R partly desensitizes C5aR1, is a new regulatory mechanism that controls receptor signals that drive the assembly and activation of the NADPH oxidase in neutrophils.

## Acknowledgement

The authors thank Linda Bergqvist for technical assistance regarding experiments.

## CRediT authorship contribution statement

**Zahra Khan**: Methodology, Formal analysis, Investigation, Visualization, Writing – Original draft

**Neele K. Levin:** Methodology, Formal analysis, Investigation, Visualization, Writing – Review and Editing

**Claes Dahlgren**: Conceptualization, Methodology, Validation, Writing – Original draft, Supervision.

**Martina Sundqvist**: Conceptualization, Methodology, Validation, Formal analysis, Investigation, Writing – Review and Editing, Visualization, Supervision, Funding acquisition. **Fariha Khan:** Supervision, Writing – Review and Editing.

**Huamei Forsman**: Conceptualization, Methodology, Validation, Investigation, Writing – Review and Editing, Supervision, Funding acquisition.

**Lena Björkman:** Conceptualization, Methodology, Validation, Formal analysis, Investigation, Writing – Review and Editing, Visualization, Supervision, Funding acquisition.

## Abbreviations

BSA: bovine serum albumin
CATPB: FFA2R antagonist - (S)-3-[2-(3-chlorophenyl) acetamido]-4-[4-(trifluoromethyl)phenyl]butanoic acid
C5aR1: the receptor that recognizes the complement component C5a
Cmp58: FFA2R allosteric modulator - ((*S*)-2-(4- chlorophenyl)-3,3-dimethyl-*N*-(5-phenylthiazol-2-yl)butanamide
FFA2R: free fatty acid receptor 2
GPCRs: G protein-coupled receptors
HRP: horseradish peroxidase
IP3: inositol- l,4,5-*tris*-phosphate
KRG: Krebs-Ringer glucose buffer
O2−: superoxide anion
PAM: positive allosteric modulator
PIP2: phosphatidyl-inositol-4,5-*bis*-phosphate
PLC: phospholipase C
ROS: reactive oxygen species
SCFAs: short chain fatty acids
TNF: tumor necrosis factor
[Ca^2+^]i: intracellular concentration of free calcium ions (Ca^2+^)

